# Survival of Thermostable Φ6 Genotypes in Acidic, Basic, and UV Environments

**DOI:** 10.1101/2025.09.24.678363

**Authors:** Aruna G. Nayagam, Elizabeth A. Wyatt, Parnian Pour Bahrami, Sonia Singhal

## Abstract

Mutations can facilitate viral resistance to a single extreme environment. But it is unknown how often these mutations confer resistance to other extreme environments to which the viruses have not been previously exposed. Here, we focus on how prior adaptation to thermal stress affects viral survival during exposure to acidic, basic, and UV conditions. We exposed four genotypes (3 single mutants and an unmutated ancestor) of the bacteriophage Φ6 with varying thermostability (low, high, or ancestral) to acid (pH 4) for 2 minutes, to base (pH 11) for 16 hours, or to UV radiation for 1 minute. We hypothesized the most thermostable genotype would also have higher survival in pH and UV stress. Unexpectedly, the least thermostable genotype demonstrated the highest survival in acidic, basic, and UV conditions, while the most thermostable genotype tended to show the lower survival in these environments. These results suggest that mutations that increase phage thermostability may not always increase pH or UV stability. Rather, molecular mechanisms of stability may depend on the environment. Investigating the survival of Φ6 in other adverse environments could provide further information on molecular mechanisms of stability and viral survival strategies.

## Introduction

Although viruses require a host to replicate, many viruses must also be able to withstand conditions in the external environment during transmission between hosts. Viruses that are more stable outside a host are more likely to maintain their structure during transmission events, improving their ability to infect their next host cell. Both the specific environmental conditions and the genotype of the virus [1, 2] play a role in its stability outside a host. Particularly for viruses that rely on indirect transmission [3-5] to spread disease, mutations that favor their stability in adverse external environments increase their chances of transmission and may contribute to viral outbreaks.

Many researchers have evaluated the survival (or inactivation) of viral particles across a range of quotidian and stressful conditions, from various surfaces to salt stress to extreme pH. For example, variola virus can be isolated from a room-temperature swab after one year [6], and some Coronaviruses, including SARS-CoV-2, can survive for up to 28 days on surfaces such as banknotes, vinyl, steel, and glass [7]. Rodent-borne Puumala virus can survive in mouse bedding between infections [8]. Influenza A virus can remain infective 207 days in 17°C water, and up to 102 days in 28°C water, before successfully transmitting to waterfowl [9]. Murine norovirus is relatively resistant to pH, while bacteriophage MS2 is relatively resistant to high salinity [10]. Other viruses, while comparatively less stable, may still maintain their structure outside a host long enough to allow a transmission event. Some influenza viruses, for example, remain viable for 48 hours on stainless steel, plastic, tissues and magazines, and respiratory syncytial virus (RSV) is viable on countertops for around 8 hours [11]. The exact conditions also matter for viral survival. For example, some cold and flu viruses decrease their stability in warm, humid conditions [12]. Chloride salts inactivate poliovirus at pH 3 but are less effective at pH 4.5-7.0 [13]. Lin et al. [14] examined the stability of bacteriophages MS2 and Φ6 in water droplets and found that chemical composition of each droplet, including pH, protein, and other environmental factors such as humidity, was crucial for viral stability. Similarly, the stability of Φ6 in water decreases with temperature [15].

The genotype of a virus can also be consequential to its stability in an external environment. Many studies have recorded the effects of mutations on the ability of viruses to withstand high temperatures (e.g., [16-24]. A mutation in influenza A virus improves its resistance to cold water during fecal-oral transmission between avian hosts [9]. A substitution in N terminal of the VP1 capsid protein increases the acid resistance of foot-and-mouth disease virus [25]. These examples demonstrate how viral genetics can increase resilience to environmental stress.

Although several studies have examined the pleiotropic effects of viral mutations across host environments (e.g., [26-30]), relatively few have evaluated the pleiotropic effects of single mutations across different external environments. (For one exception, see [23], where a poliovirus mutant was evaluated for both thermostability and bleach resistance.) Such information could be important both for generating epidemiological predictions as environments change due to climate change, and for establishing molecular mechanisms of viral stability.

We address pleiotropic effects of mutations using Φ6 Cystovirus, a well-studied bacteriophage (phage) that infects the bacterium *Pseudomonas syringae* pathovar *phaseolicola*. Φ6 is a lytic, double stranded RNA bacteriophage in the Cystoviridae family and has a genome composed of three segments (L, M, and S). Unlike most phages, but similar to most animal viruses, Φ6 is enveloped with a phospholipid membrane [31], and it shares structural and assembly features with Reoviruses [32, 33], making it a valuable non-pathogenic model.

A prior study by Singhal et al. [21] evolved populations of Φ6 for greater thermostability by exposing them to periodic, high-temperature heat shocks over 32 transfers (approximately 100 viral generations). Each transfer in the experiment followed a consistent cycle: phage lysates were first exposed to a five-minute heat shock up to 50°C, and the surviving phages then infected naïve cultures of *Pseudomonas phaseolicola* at 25°C. Because viruses were exposed to heat stress as inert particles, when protein activity of their attachment and replication machinery was inactive, the transfer regime promoted the evolution of heat resistance through enhanced structural integrity, or greater protein stability. From the evolved populations, Singhal et al. identified 10 unique mutations in the P5 lysis protein [21], which is found in the nucleocapsid outer shell of Φ6 and has a dual function for both early infection (penetration of the host cell wall by the viral core) and late infection (lysis of the host to release progeny phage, [34, 35]). Singhal et al. characterized 6 of these P5 mutations for their effects on viral thermostability [21].

We chose four genotypes from Singhal et al. (2017), representing varying degrees of tolerance to thermal stress (Figure 1, [21]), to examine the pleiotropic effects of these mutations in non-thermal environments. The locations of these mutations in the P5 protein are shown in Figure 2. The ancestor acts as a wild type genotype and has no mutations. Genotype P5 W219^*^ has a mutation in the C-terminal tail of the P5 lysin, and has a thermostability similar to the ancestor’s. Genotype P5 R124G has a mutation on the exterior of the lysin protein, and has reduced thermostability compared to the ancestor. Genotype P5 V207F has a mutation on helix a8, facing the hydrophobic core of the protein close to the C terminal region. This mutation confers high heat resistance [19, 21], has been demonstrated to increase the stability of the P5 protein [18] and is commonly found when Φ6 is exposed to high temperatures [16,18, 19, 21]. Apart from P5 V207F, the molecular mechanisms that modulate thermostability of these genotypes have not been investigated.

**Figure 1:**
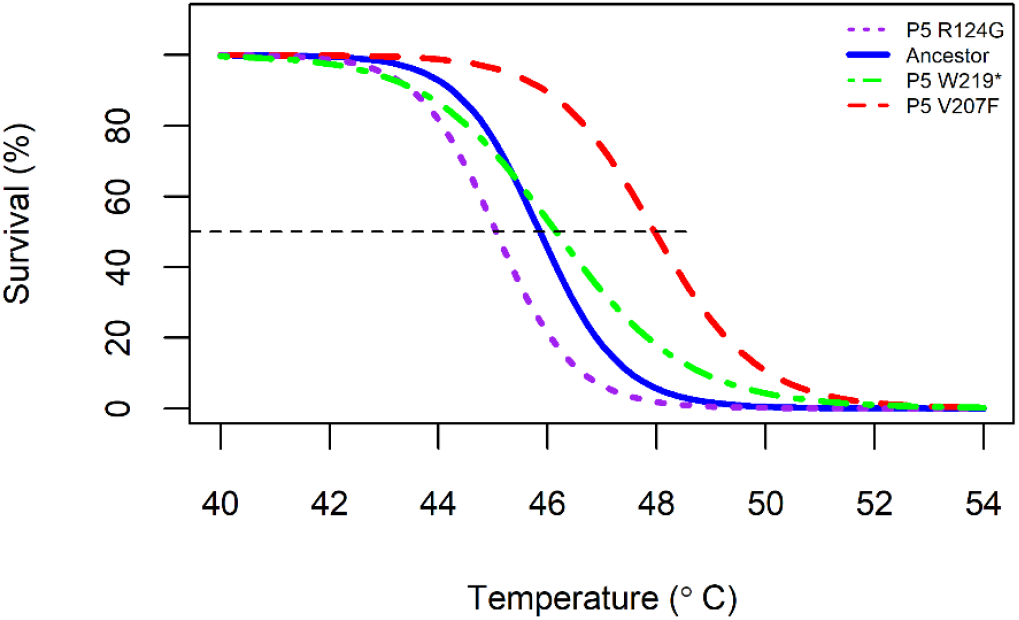
Thermal kill curves for the four Φ6 genotypes used in in this study. The dotted line indicates the inflection point of the curves, where 50% of viruses of that genotype survive heat stress, and represents genotype thermostability. The ancestor is unmutated and is used as the control; P5 R124G is the least thermostable genotype; P5 W219^*^ has thermostability similar to the ancestor’s; and P5 V207F is the most thermostable genotype. Data from [21].

**Figure 2:**
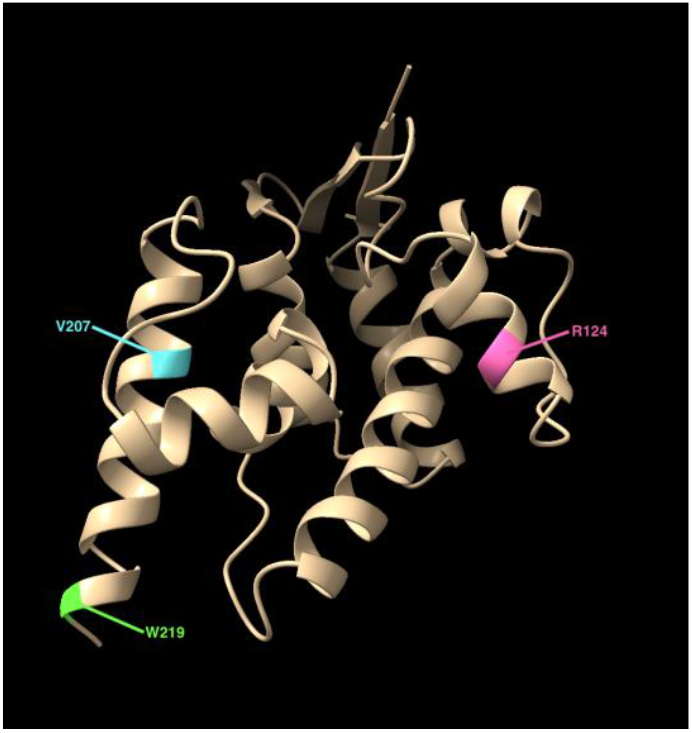
Ribbon diagram of the Φ6 P5 lysis protein rendered in UCSF Chimera [36], highlighting the positions of mutations. Residue R124 is shown in pink, V207 in cyan, and W219 in green. The structure was visualized using PDB entry 4DQ5.

Here, we examine whether these thermostabilizing and non-thermostabilizing mutations also confer resistance to three other environmental stressors: exposure to acid (pH 4), base (pH 11, and UV radiation. As in Singhal et al. [21], we isolate stability effects by exposing inert viral particles to the stressful environment, and then assaying their survival with plating assays under standard growth conditions. We hypothesize that there is a positive correlation between thermostability and pH or UV stability: the phage genotype with highest thermostability (P5 V207F) will also have the highest survival in acidic, basic and UV environments. Our research has wider implications for understanding how viruses evolve in changing environments, the effects of single mutations on viral proteins, and the molecular mechanisms of protein stability in viruses.

## Materials and Methods

### Genotypes and culture conditions

We use four Φ6 genotypes (Ancestor, P5 R124G, P5 W219^*^, P5 V207F) that differ from each other by one nucleotide [21]. These genotypes display different levels of resistance to high temperatures (Figure 1).

*Pseudomonas syringae* pathovar *phaseolicola* (derivative of ATCC #21781) was used as a host. Bacteria were grown from isolated colonies in LC broth (Luria-Bertani broth at pH 7.5) overnight at 25°C, with shaking at 210 rpm.

To make working stocks of phage, 100 µL of a dilution of phages and 200 µL of overnight bacterial culture were added to 3 mL of molten LC 0.7% soft agar and overlaid on a LC 1.5% agar base. After 24 hours of incubation at 25°C, plaques were collected from plates that showed complete, confluent lysis of bacterial cells and transferred into 3 mL of LC broth. The solution was vortexed, and viruses were separated from the agar by centrifugation for 7 min at 3000 rpm, followed by filtration through a 0.22 µm cellulose acetate filter. The resulting viral lysates were stored at 4°C for working use.

Phage concentrations were determined by plating serially diluted samples on a *P. phaseolicola* lawn using the agar overlay method described above and incubating overnight at 25°C. To avoid the effects of varying viral concentrations on viral survival, all viral lysates were normalized to a target concentration of 2.17 x 10^9^ plaque-forming units (pfu)/mL for experimental use.

Acid and base exposure occurred in LC medium, adjusted to pH 4 using citric acid or pH 11 with sodium hydroxide. UV exposure occurred in SM phage buffer [37], which is optically transparent.

### Determination of bacterial survival in acidic and basic environments

To ensure that phage death in different pH environments was due to their inability to survive pH stress, and not due to bacterial death at extreme pH values, we first confirmed that the *P. phaseolicola* host could survive in the presence of small volumes of acidic and basic media to allow phage propagation. Volumes from 200 µL to 500 µL of overnight bacterial culture, along with 100 µL of pH-adjusted medium (the volume used when plating phages), were combined in 3 ml of LC 0.7% soft agar, overlaid on a LC 1.5% agar plate, and incubated overnight at 25°C. Here, we plated bacterial samples in the absence of phages to confirm a thick lawn of bacterial growth. We observed a thick lawn when using 500 µL of culture. This served as the culture volume for agar overlay plating for the remainder of the study.

### Time series assay

A time series assay conducted with the ancestral genotype was used to determine the exposure time of phages to each stress. The initial concentration of phages was determined by plating a stock lysate onto a *P. phaseolicola* host lawn. A phage sample was exposed to the acidic medium for up to 5 minutes, with samples taken every 1 minute, and to the basic medium for up to 24 hours, with samples taken after 0 minutes, 30 minutes, 60 minutes, 90 minutes, and 24 hours. Similarly, phages were exposed to UV-C radiation using a UV illuminator at 254 nm, with samples taken every 1 minute. After each exposure, phages were plated on a *P. phaseolicola* lawn to determine the number of surviving viruses. The time point at which approximately 10% of the phages survived was used as the exposure time.

### pH and UV survival assay

We determined the survival of the four viral genotypes (3 mutants and the unmutated ancestor) in acid, in base, and under UV irradiation using the time determined from the time series assay. Initial phage titers for each genotype were confirmed using the agar overlay method before exposing the phages to acid, base, or UV for 2 minutes, 16 hours, or 1 minute, respectively. The phage titer after exposure was again determined by agar overlay plating. A neutral (pH 7) or non-irradiated environment served as the control.

### Statistical analysis

Survival differences among genotypes and pH environments were analyzed with ANOVA and Tukey’s Post Hoc test. Statistical analysis was performed in R studio version 2024.04.1+748.

## Results

We examined survival of the ancestor (unmutated) Φ6 genotype and three genotypes with a single-base substitution that affected the virus’s thermostability. The ancestor has no mutations and is used as the control, P5 R124G is the least thermostable genotype, P5 W219^*^ has thermostability similar to the ancestor’s, and finally P5 V207F is the most thermostable genotype (Figure 1).

### Determination of exposure time for acid, base and UV stress

The ancestor phage was exposed to LC medium adjusted to pH 4 or 11 to determine the exposure time that would result in a decrease in survival to below 10%. In the acidic environment, the ancestor showed a rapid decrease in survival; survival dropped well below 10% (to 0.001%) within the first two minutes. Survival further decreased to 0.0005% after 3 minutes of exposure, and no surviving phages were detected after 4 minutes of exposure (Figure 3). We therefore chose 2 minutes as a suitable exposure time for the acidic environment.

**Figure 3:**
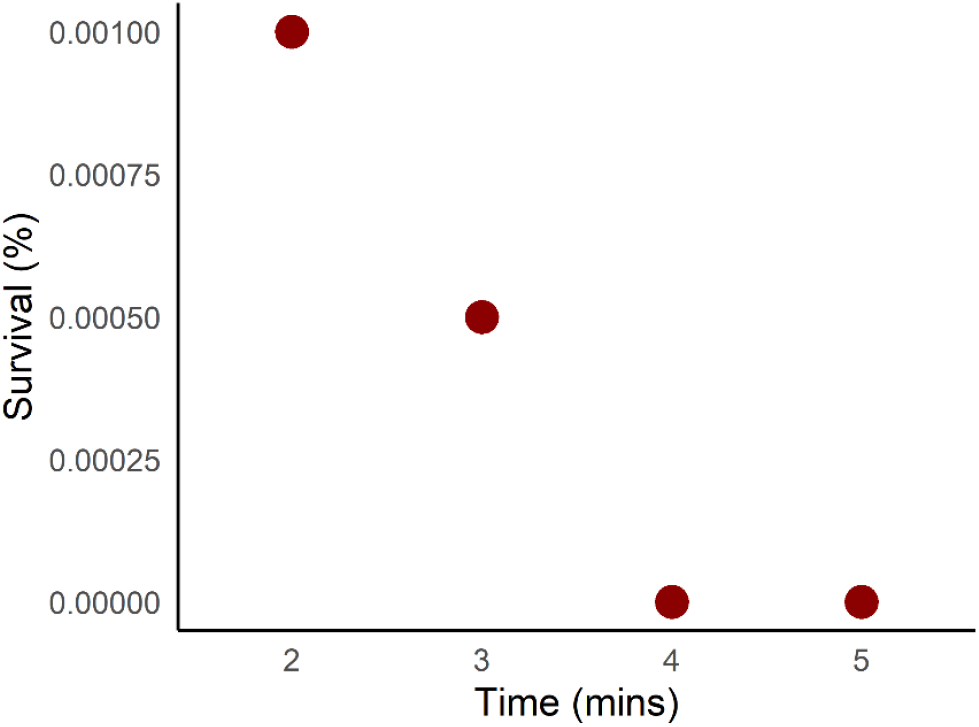
Survival of ancestor phage when exposed to pH 4 between 2-5 minutes. Each point represents the average of 2 replicates.

In the basic environment the survival showed a monotonic decline over 24 hours. We fit a trendline to the data and solved the equation of this line for Y = 10% survival, resulting in an exposure time of 16 hours. We used this time as the exposure time for the basic environment (Figure 4). Under UV stress, the ancestor phage showed 0.8% survival at 1 minute (Figure 5), and no survival was detected at later time points. We therefore used an exposure time of 1 minute for UV survival assay.

**Figure 4:**
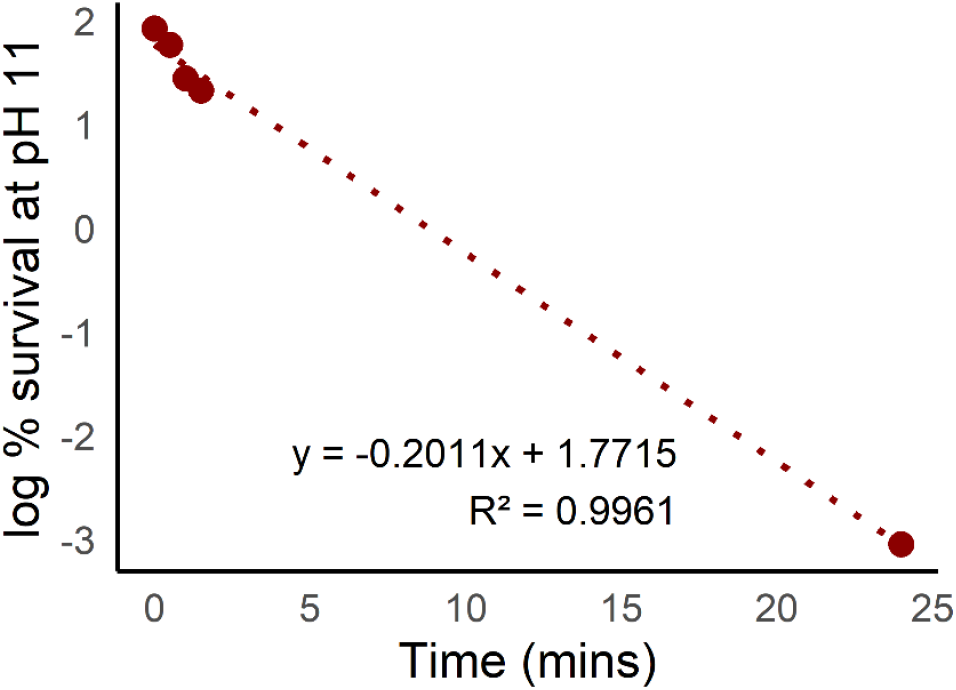
Survival of ancestor phage when exposed to pH 11 for 0, 30, 60 and 90 minutes, and 24 hours. Each point represents the average of 2 replicates. Dotted line indicates the regression line of the log-transformed data.

**Figure 5:**
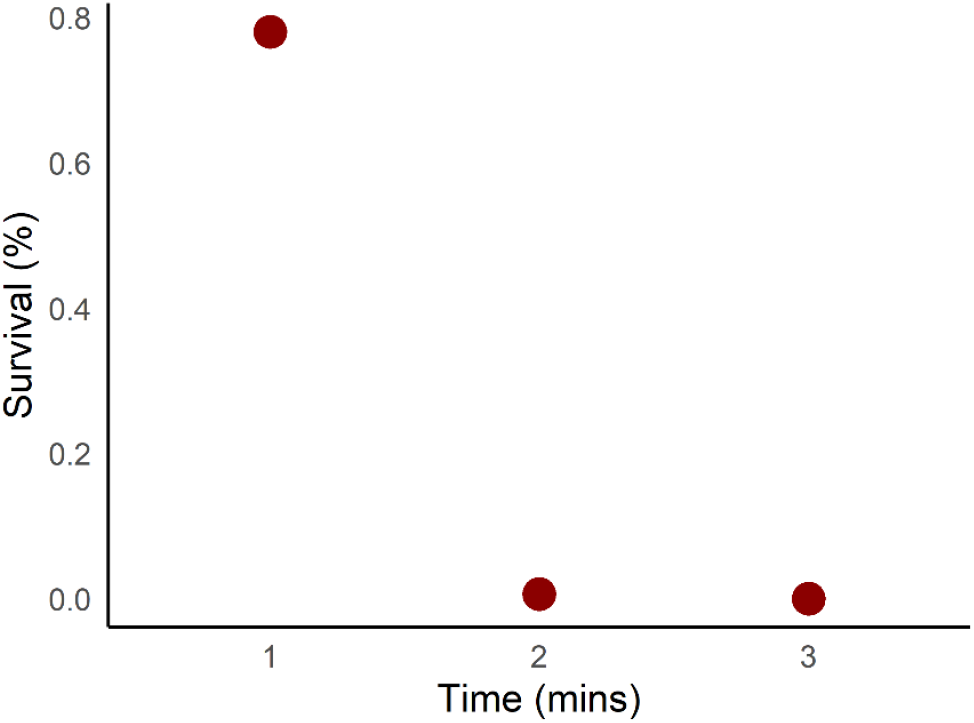
Survival of ancestor phage when exposed to UV irradiation between 1-3 minutes. Each point represents the average of 2 replicates.

### Genotype with low thermostability shows greatest survival in acid and base stress

All Φ6 genotypes, including the unmutated ancestor phage (n=8), showed 100% survival in the control pH 7 environment. In comparison to the control environment, survival of the ancestor decreased significantly in pH 4 medium (2.8% survival) and in pH 11 medium (2.1% survival, Tukey’s post-hoc test, p << 0.001; Figure 6). Surprisingly, P5 R124G (the least thermostable genotype) showed the highest survival in both acidic (12.8% survival, Tukey’s post-hoc test, p << 0.001) and basic (11.4% survival, p << 0.001) conditions. In contrast, P5 V207F (the most thermostable genotype) tended to show the lowest survival in acid (0.1% survival) or base (1.4 % survival, Figure 6), although this difference was not significantly different from the ancestor.

**Figure 6:**
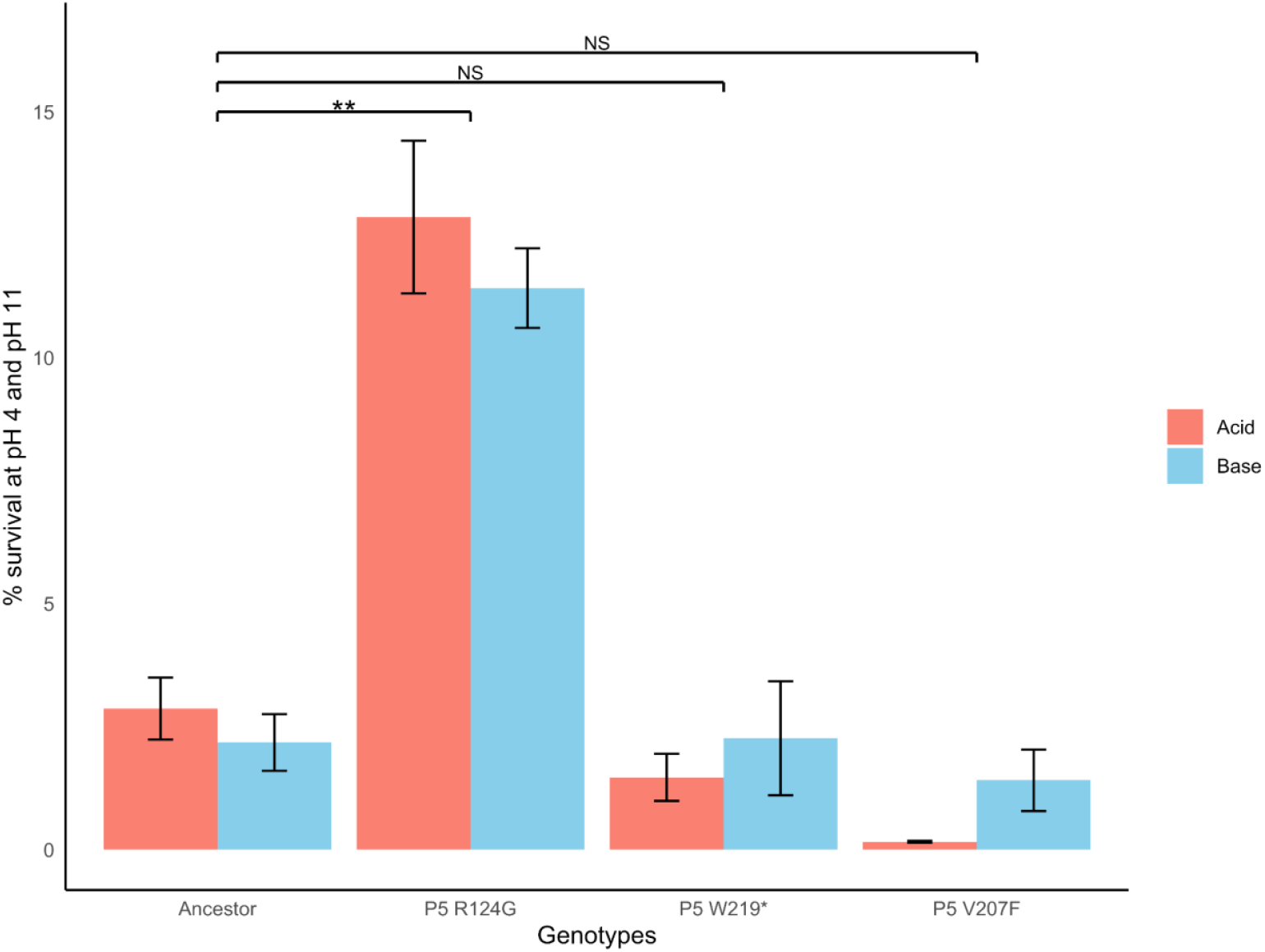
Survival of thermostable genotypes at pH 4 and pH 11. The phages were exposed to acid for two minutes and base for 16 hours and plated immediately in *P*.*Phaselicola* lawn. Each bar represents the average of 8 replicates; error bars denote standard deviation. Only P5 R124G survived significantly better than the ancestor (ANOVA, p << 0.01).

### Genotype with low thermostability shows greatest survival under UV exposure

Phages exposed to UV were compared to phages that did not experience UV exposure. In the control environment (SMB buffer with no UV irradiation) there was 100% survival. The average survival of all four genotypes after 1 minute of UV exposure was 8.3%. P5 R124G (the least thermostable genotype) showed the highest survival (15.3% survival, p<<0.01). P5 W219^*^ (genotype closest to the wild type in thermostability) tended to have the lowest survival (5.5% survival), although the difference was not statistically significant in comparison to the ancestor (Figure 7).

**Figure 7:**
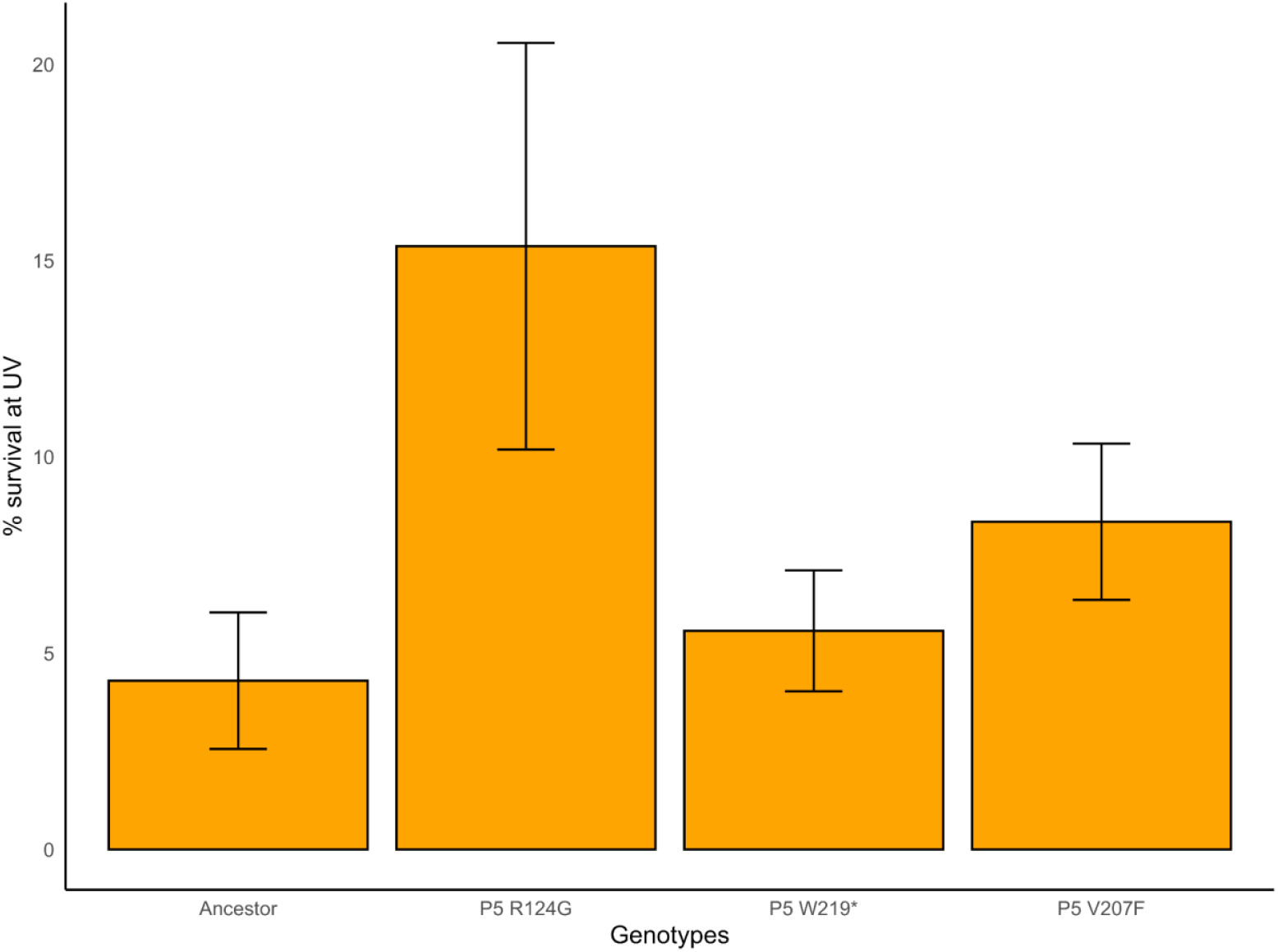
Survival of thermostable genotypes after 1 minute of exposure to UV radiation. Each bar represents the average of 8 replicates; error bars denote standard deviation. Only P5 R124G survived significantly better than the ancestor (ANOVA, p<< 0.01).

## Discussion

By evaluating the survival of four Φ6 genotypes that varied in their thermostability, we demonstrate that a single mutation in one protein can also affect the ability of phages to survive in stressful abiotic pH and UV conditions. In acidic and basic environments, the overall survival of the four genotypes was less than 10%, despite remaining at 100% in the neutral (pH 7) environment. The highly thermostable P5 V207F genotype tended to show the lowest survival (less than 2% survival), while the non-thermostable P5 R124G genotype had significantly higher survival (approximately 12% survival) than the ancestor in these environments. In the UV environment, P5 R124G retained the highest survival (15.3% survival). This pattern suggests that the mutations that increase thermostability may not always increase the pH or UV stability of the phages. Instead, the mechanism of protein stability may change based on the environment.

A mechanism for thermostability was previously established for P5, the protein in which our genotypes differ. P5 encodes the viral lysin, a lytic transglycosylase [38] used by Φ6 to penetrate the bacterial cell wall during entry into and exit from the host cell [18, 34]. Crystal structures of ancestral (wild type) P5, P5 with the V207F mutation, and ligand-bound P5 reveal the molecular basis of its thermostabilization: the Phe207 side chain fills a hydrophobic cavity that is unoccupied in the wild-type. The mutation raises the melting point of the protein, as measured by circular dichroism spectrometry and differential scanning calorimetry, most likely by increasing the molecular van der Waals interactions between amino acids [18, 39].

Unlike the thermal environment, where hydrophobic interactions are often a dominant factor influencing protein stability, pH stress primarily challenges the ionization states of amino acid side chains. The overall charge state of a protein is shaped by how its ionizable residues, such as lysine, arginine, aspartate, or glutamate, respond to the surrounding pH, which can have major effects on conformation, solubility, and stability of the protein [40, 41]. Protonation or deprotonation events can disrupt local hydrogen bonding networks and electrostatics, ultimately destabilizing the protein [42, 43]. Arginine, for example, carries a guanidinium group that typically remains positively charged under physiological conditions; yet shifts in its microenvironment may alter ionization and affect local structure [44]. In contrast, glycine lacks an ionizable side chain, consisting only of hydrogens, and therefore remains unaffected by pH changes. In this way, residues such as glycine may promote greater pH stability, whereas ionizable residues such as arginine may introduce conformational instability under changing pH conditions.

Ionization states of amino acids may also alter protein conformation. For instance, a change from arginine to glycine (e.g., R124G) eliminates protonation/deprotonation events that could alter conformations, while also removing steric bulk in sensitive structural regions. Surface views of the P5 lysin show that Arg124 fills a prominent surface cavity in the wild-type, whereas the substitution to glycine produces a distinct gap, highlighting both the loss of charge and altered amino acid packing at this site (Figure 8). Glycine’s non-ionizable nature and small size prevents local pH-driven destabilization while still supporting conformational flexibility. This concept aligns with previous studies showing that altering specific ionizable residues can markedly shift the pH stability profile of proteins [45-47].

**Figure 8:**
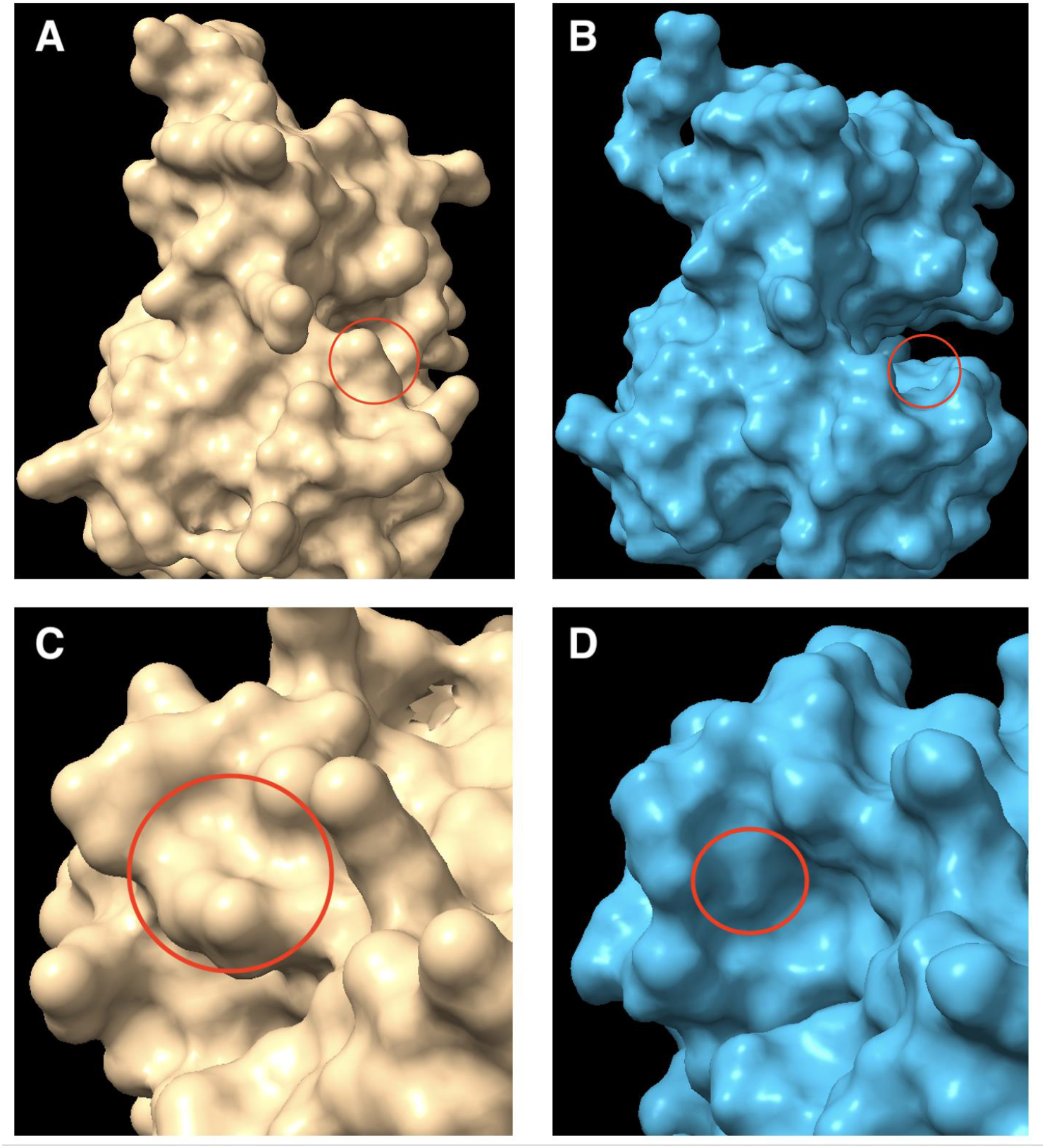
Surface representations of the Φ6 P5 lysis protein, highlighting the structural impact of the R124G mutation. The wild-type protein is beige, with arginine (R124) circled in red (A, C). The mutant protein is blue, with glycine (G124) circled in red (B, D). Views from different orientations show that arginine fills a prominent surface cavity, while substitution to glycine generates a marked gap at position 124. Structures were predicted using AlphaFold [48] and visualized with UCSF Chimera [36].

Ogbunugafor et al. [49] previously examined stability of Φ6 under UV radiation. They exposed two Φ6 genotypes to UV-C (254 nm) radiation for between 5 and 15 minutes. The specific genotypes originated from an evolutionary study [50, 51] where Φ6 was evolved at either high or low multiplicity of infection (ratio of viral particles: bacterial cells), and thus contained different mutations than the ones in our study. Ogbunugafor et al found no statistically significant differences in survival between their two genotypes of Φ6 [49]. Our study suggests, however, that certain amino acid substitutions can influence virion stability under UV irradiation. Our least thermostable strain, R124G, showed the highest survival (15.3% survival) with UV exposure, while the strain with thermostability closest to the ancestor, W219^*^, showed the lowest survival (5.5% survival). Aromatic amino acids with benzene side chains (tryptophan, tyrosine and phenylalanine) can absorb UV radiation from 265 nm, making them more vulnerable to UV damage. Upon absorption, they undergo electronic excitation, leading to photooxidation, generation of reactive oxygen species [52], and possibly chemical modifications [53] of the side chains—ultimately causing protein damage or functional loss [54].

Our work highlights that single mutations can result in differences in survival across multiple environments. We demonstrate that the molecular mechanism for stability in thermal, pH, and UV stress depends on how the viral proteins interact with the external stressor, which in turn depends on the location of the mutation within the protein, the size and structure of the amino acid, and the molecular mechanism of resistance to the stressor. Further evaluation of the P5 wild type and mutated P5 proteins would allow us to confirm our hypothesized molecular mechanisms of pH and UV stability. Testing the survival of these genotypes in other extreme environments that viruses might encounter outside a host cell could also provide more information about viral survival strategies.

## Acknowledgements

This research was supported by the National Institute of General Medical Sciences (Grant R16GM146706 to SS) and Research and Innovation Student Research, Scholarship, and Creative Activities Fellowship from San Jose State University (Award # 24-SRF-08-087 to AGN).

## Author Contributions

Project administration: A.G.N. Conceptualization, Methodology, Visualization: A.G.N., B.W., S.S. Investigation, Data curation, Formal analysis, Validation: A.G.N., B.W. Funding acquisition: A.G.N., S.S. Writing—original draft: A.G.N., B.W., P.P.B. Writing—review & editing: A.G.N., B.W., P.P.B., S.S. Resources, Supervision: S.S. All authors have read and agreed to the published version of the manuscript.

## Conflicts of Interest

The authors declare no conflicts of interest.

## Notes

### Competing Interest Statement

The authors have declared no competing interest.

